# Selection mode changes the fate of nearly neutral deleterious mutations

**DOI:** 10.64898/2026.03.04.709298

**Authors:** Thomas Birley, Cock van Oosterhout

## Abstract

Nearly neutral theory predicts that deleterious alleles can behave approximately neutrally when selection is weak relative to genetic drift. For codominant mutations, the conventional nearly neutral threshold is |*s*| ≈ 1/(2*Ne*); thus, when *Ne* = 500, a mutation with |*s*| = 0.001 lies at this threshold. For strongly recessive mutations, however, newly arisen copies occur mainly in heterozygotes, so selection is much weaker while they are rare. We used forward-time simulations to test whether competitive soft selection can purge strongly recessive deleterious mutations that behave as nearly neutral under hard selection. The focal mutations had *s* = -0.001 and *h* = 0.01, placing their effect when rare about 50-fold below the corresponding nearly neutral threshold. Under hard selection, these mutations behaved almost neutrally, continuing to accumulate, with some reaching fixation. Under rank-based soft selection, accumulation was strongly suppressed; at high fecundity, mutations remained at low frequency and none reached fixation. Purging strengthened as more candidates competed for a fixed number of recruitment opportunities. A pairwise tournament produced the same qualitative but slightly weaker pattern. Thus, competitive soft selection can purge rare, recessive deleterious mutations whose effects fall well below the nearly neutral threshold under hard selection. By acting on aggregate multilocus fitness differences, relative competition may reduce genetic load and susceptibility to inbreeding depression. These findings are also relevant to genic capture, whereby many weak effects of deleterious mutations contribute to genome-wide genetic quality that can be exposed through competition and sexual selection.

## 1. Introduction

Nearly neutral theory describes how weakly deleterious mutations can behave effectively neutrally when selection is weak relative to genetic drift [1–4]. A common rule of thumb places the nearly neutral threshold near |*s*| ≈ 1/(2*Ne*), but the relevant selective effect depends on how *s* is defined and on its dominance coefficient, *h*. A newly arisen diploid deleterious allele is usually carried in a heterozygote, so selection initially acts through *hs*. For codominant mutations with *h* = 0.5, the formulations |*s*| ≈ 1/(2*Ne*) and |*hs*| ≈ 1/(4*Ne*) are equivalent. With strong recessivity, a deleterious allele can remain weakly exposed to selection while rare and therefore drift to higher frequency, contributing to masked genetic load (Ohta 1973; Charlesworth 2009; Bertorelle et al. 2022). As homozygosity increases, this hidden load is exposed, increasing susceptibility to inbreeding depression (Dussex et al. 2023).

The mode of selection can further alter how genotype-level fitness differences are converted into survival or reproduction. Wallace distinguished hard selection from soft selection [5]. Under hard selection, success depends on absolute fitness, irrespective of the fitness of other individuals. Under soft selection, a fixed number of ecological opportunities are allocated according to relative performance [5]. In a rank-based competitive setting, increasing the number of candidates competing for the same number of opportunities increases the intensity of filtering because success depends on relative performance within the candidate pool [5–7]. Populations with the same number of breeders can therefore experience similar genetic drift while differing substantially in the strength of purifying selection.

Relative competition is common during recruitment, territorial competition and mating. Sexual-selection theory has long recognised that reproductive success can expose differences in genetic quality [8–10], and genic capture provides a mechanism by which condition-dependent traits integrate effects across many loci [11]. Genomic data increasingly support this connection. Stronger sexual selection is associated with more efficient purifying selection across birds [12], genomic deleterious load predicts male mating success in wild black grouse [13], and experimental evolution in *Tribolium* shows that stronger sexual selection can reduce deleterious variation and subsequent extinction risk [14]. These studies make it important to distinguish the fitness effect of a mutation from the ecological process that converts that effect into realised reproductive or survival differences.

Here we use forward-time simulations comparing hard selection with an idealised competitive form of soft selection. The focal mutations were deleterious (*s* = -0.001) and strongly recessive (*h* = 0.01). At *Ne* = 500, their homozygous effect therefore lies at the conventional nearly neutral threshold, whereas a newly arisen copy occurs in a heterozygote and experiences a much weaker effect, *hs* = -1 × 10□□. We test whether competitive soft selection can purge these mutations even when they are expected to behave as nearly neutral under hard viability selection. We also test whether purging becomes stronger as more candidates compete for a fixed number of recruitment opportunities, and whether the same qualitative pattern occurs under pairwise tournament competition.

## 2. Simulations

We used forward-time individual-based simulations in SLiM 4.3.0 [15]. Populations were diploid, hermaphroditic and non-overlapping, with carrying capacity *K* = 500. The simulated genome contained 20 independently assorting autosomes, with the genome architecture, recombination rates and mutation scheme described in the Electronic Supplementary Material. Each parameter combination was replicated 100 times and followed for 5,000 generations.

The main comparison used deleterious mutations with homozygous selection coefficient *s* = -0.001 and dominance coefficient *h* = 0.01. A newly arisen copy therefore had a heterozygous effect *hs* = -1 × 10□□. Realised effective population size (*Ne*) was estimated using four complementary approaches: nucleotide diversity, temporal allele-frequency change, mean pairwise coalescence time, and variance in realised reproductive success. The first two used neutral marker mutations introduced during the burn-in, while the coalescent estimate was obtained from SLiM tree sequences. Full details are provided in the Electronic Supplementary Material. Across estimators and treatments, *Ne* ranged from approximately 475 to 564 (Table S1). Thus, at *Ne* ≈ 500, *s* = -0.001 lies at the conventional codominant nearly neutral threshold, whereas selection on a newly arisen copy is 50-fold weaker than for a codominant mutation with the same *s*.

Under hard selection, each candidate survived with probability equal to its genotype-level fitness. If more than *K* individuals survived, random density regulation reduced the population to *K*. Under rank-based soft selection, candidates received a competitive score equal to 0.5*w*□ + *U*(0,0.5), where *w*□ is the genotype-level fitness of individual *i*. The *K* highest-ranking candidates survived. Fecundity generated either 5*K* or 50*K* candidates for the same *K* recruitment opportunities, representing low- and high-fecundity conditions. The ranking model therefore represents a strong, single episode of cohort-wide competitive filtering.

We also examined soft selection in a pairwise tournament. Candidates were repeatedly paired and the individual with the higher competitive score survived each round until *K* individuals remained. Higher fecundity therefore generated more elimination rounds before recruitment. Full implementation details are given in the Electronic Supplementary Material. We recorded mutation accumulation and the site-frequency spectrum (SFS), and compared neutral evolution, hard selection and both forms of soft selection.

## 3. Results

### Hard selection vs rank-based soft selection

Under hard selection, the focal mutations accumulated through time and followed a trajectory much closer to neutrality than to either soft-selection treatment (Figure 1a). Increasing fecundity from 5 to 50 had little qualitative effect on mutation accumulation under hard selection. This is expected because each individual’s survival probability depends on its own fitness rather than on its performance relative to other candidates.

**Figure 1.**
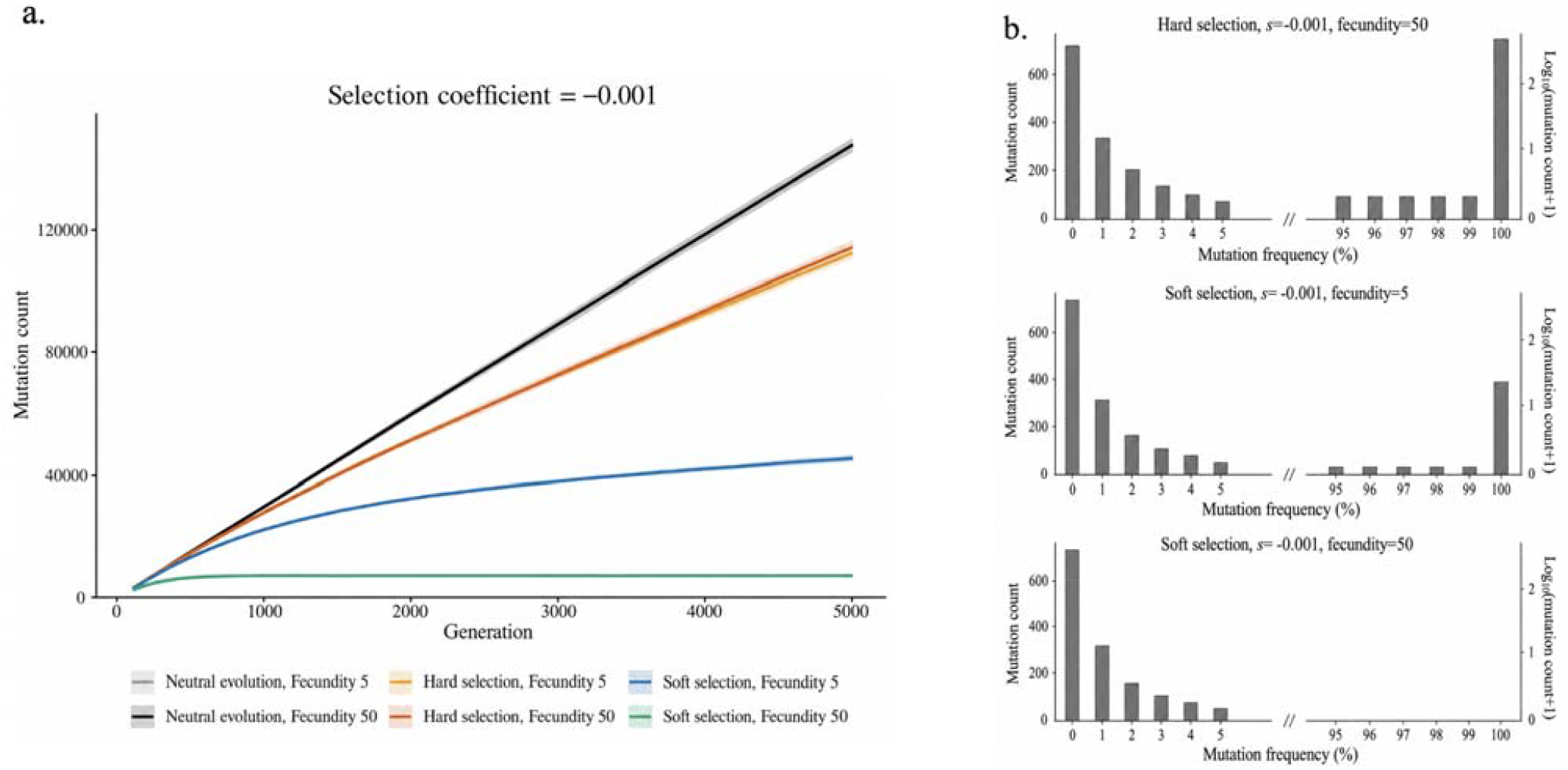
Competitive soft selection purges deleterious mutations that behave as nearly neutral under hard selection. (a) Accumulation of mutations with *s* = -0.001 and *h* = 0.01 over 5,000 generations under neutral evolution, hard selection and rank-based soft selection, with fecundity set to 5 or 50. Under hard selection, mutations continued to accumulate and their trajectories were much closer to neutrality than under soft selection. Increasing fecundity had little effect under hard selection. Under soft selection, mutation accumulation was strongly reduced and purging became more efficient as more candidates competed for the same *K* recruitment opportunities. (b) Site-frequency spectra (SFS) for the same mutation class at generation 5,000. Hard selection retained a pronounced high-frequency tail, including fixed mutations. Soft selection reduced intermediate- and high-frequency variants, with the strongest effect at fecundity 50, where mutations remained concentrated at low frequency and none reached fixation.

Rank-based soft selection produced a much stronger response. At fecundity 5, mutation accumulation slowed substantially, while at fecundity 50 it levelled off at a low value after the initial generations. Purging therefore became stronger as more candidates competed for the same *K* recruitment opportunities. Mutations that behaved as nearly neutral under hard selection were thus efficiently purged under strong competitive soft selection.

The site-frequency spectra (SFS) showed the same pattern (Figure 1b). Hard selection retained a pronounced high-frequency tail, including fixed mutations. Soft selection reduced intermediate- and high-frequency variants, with the strongest effect at fecundity 50. Under high-fecundity soft selection, almost all focal mutations remained at very low frequency, and none reached fixation by generation 5,000. Competitive filtering therefore affected both the accumulation of deleterious mutations and their probability of reaching high frequency.

### Contest model: algebraic formulation

To examine how repeated pairwise assessments can amplify a persistent fitness difference, realised load can be represented on a log-fitness scale [16]. If realised load is and competitive ability is proportional to, then for two individuals A and B with loads and, the probability that A wins a single contest is

If the same pair competes independently *m* times, the number of contests won by A follows

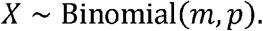

For an odd number of contests, the probability that A wins the series is

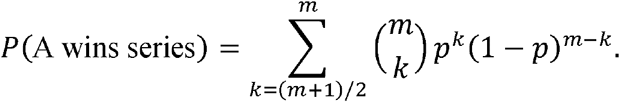

For *m* = 1, this reduces exactly to *P* = *p*. As *m* increases, repeated contests make a persistent difference in realised load increasingly likely to determine the overall winner. Figure 2 illustrates this increase in discrimination across repeated competitive assessments.

**Figure 2.**
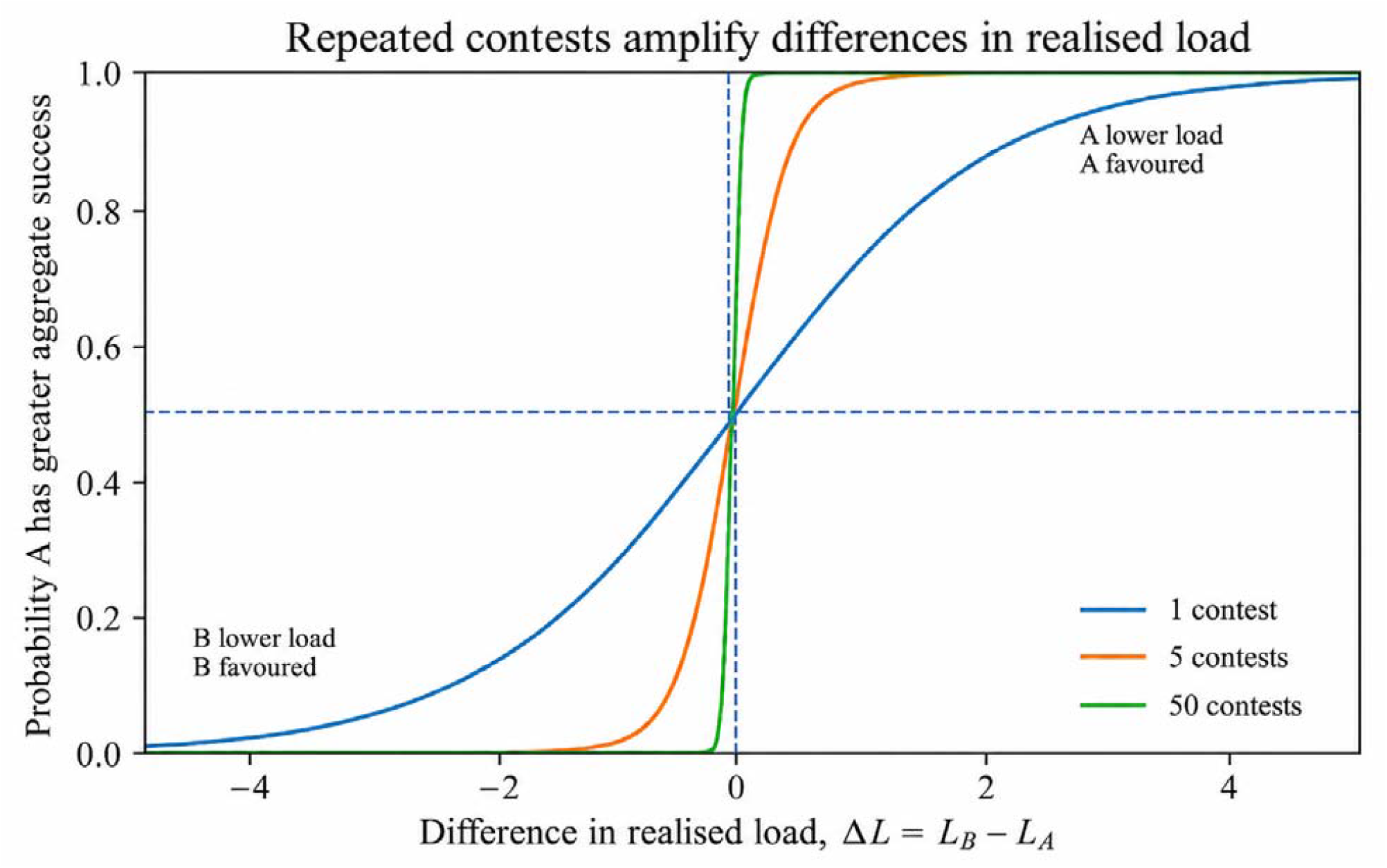
Repeated competitive assessment amplifies differences in realised load. Repeated exposure to the same persistent fitness difference increases the probability that the lower-load individual wins the contest series.

The pairwise tournament produced the same qualitative (albeit weaker) pattern as the rank-based soft selection model (Electronic Supplementary Material, Figure S2). Mutation accumulation again declined as the number of candidates increased. Realised *Ne* remained similar across selection modes (Electronic Supplementary Material, Table S1), indicating that differences in genetic drift cannot explain the contrast in mutation accumulation.

## 4. Discussion

Nearly neutral theory predicts mutation fate from the balance between selection and genetic drift [2]. Our results are consistent with this framework: competitive structure can change the strength of selection experienced by a mutation even when its underlying genotype-level effect is unchanged. This is especially relevant for recessive deleterious variants, whose effects are weakly expressed when rare. We therefore tested whether mutations that behave as nearly neutral under hard viability selection can nevertheless be purged when selection acts through relative competition. In our model, deleterious mutations (*s* = -0.001) were strongly recessive (*h* = 0.01), giving an effect of only *hs* = -1 × 10□□ when rare. Under hard selection they accumulated, with some mutations reaching fixation. Under soft selection, the same mutations accumulated much more slowly, and purging became stronger as more candidates competed for the same number of recruitment opportunities. The pairwise tournament produced the same qualitative, although weaker, pattern. Because realised *Ne* remained similar among treatments, this contrast reflects differences in selection rather than differences in genetic drift.

Our results can be understood by considering how fitness determines recruitment success. Under hard selection, each candidate experiences the same absolute survival function. Increasing fecundity exposes more individuals to selection, but it does not change the fitness function itself. In the rank-based model, recruitment depends on relative performance. At higher fecundity, breeding individuals are recruited from a much larger pool and therefore from a more extreme upper tail of the competitive-score distribution. The selected individuals consequently represent a smaller fraction of the highest-performing candidates in the population. A small genotype-dependent shift in score can consequently have a much larger effect on recruitment when competition is intense.

The rank-based model represents the strongest form of competitive filtering and provides an upper bound on this effect. The pairwise tournament is less discriminating because individuals are compared locally rather than ranked across the entire cohort. Still, even local contests can strengthen the efficiency of purging. The algebraic contest model illustrates the same principle at the level of repeated assessment. A small difference in realised load has only a modest effect on the outcome of one contest, but repeated independent contests make the lower-load individual increasingly likely to win the series. Competitive structure therefore determines how strongly small fitness differences are translated into differences in realised success.

This is particularly relevant for variants around the nearly neutral threshold. In our simulations, selection acted on total multilocus fitness, so many individually small effects contributed to differences among genetic backgrounds. Competitive filtering acted on those aggregate differences and reduced the probability that individual deleterious mutations progressed to high frequency. Strong relative competition may therefore limit the build-up of recessive variation that can later be exposed by homozygosity and contribute to inbreeding depression.

This mechanism is relevant to genic capture, whereby condition-dependent traits integrate genetic effects across many loci and thereby expose genome-wide genetic quality to selection [11]. This allows genome-wide deleterious variation to influence competitive or reproductive performance. Mate choice, lek attendance, territorial competition and other condition-dependent behaviours can therefore expose small differences in genetic quality to relative selection [8–14]. Recent genomic results are consistent with this idea: stronger sexual selection is associated with more efficient purifying selection across birds [12], genomic deleterious load predicts male mating success in black grouse [13], and experimental evolution in *Tribolium* shows that stronger sexual selection can reduce deleterious variation and subsequent extinction risk [14]. Our models provide a simple mechanism through which such effects can arise when reproductive success depends on relative performance.

Sexual selection is only one possible setting for this process. Competition for territories, breeding vacancies, parental resources or recruitment opportunities can generate the same type of relative filtering. The important quantity is the relationship between the number of candidates and the number of opportunities, together with how accurately competitive performance reflects underlying genetic fitness. Environmental variation, the number of rivals, repeated assessment and stochastic contest outcomes will all weaken or strengthen that relationship. The contrast between the global ranking and pairwise tournament models illustrates how strongly the outcome can depend on competitive structure.

The simulations show this in a simplified form, using one class of strongly recessive deleterious mutations, fixed population size and simplified competitive rules. Natural populations contain distributions of fitness effects and dominance coefficients, linkage among loci, environmental variation and competition operating at several stages of the life cycle. These factors will determine the quantitative strength of competitive purging. The present result is therefore a prediction about mechanism: for a given *Ne* and mutational effect, increasing the extent to which recruitment depends on relative genetic performance should reduce the persistence of weakly expressed deleterious variation.

Because the same underlying fitness effect can have different evolutionary consequences under different forms and intensities of competition, a useful next step would be to place these selection regimes on a common quantitative scale. Such an approach would allow direct comparison of how strongly different ecological settings weaken or strengthen purifying selection relative to hard selection. This would also improve the interpretation of genomic proxies of genetic load, because the same deleterious variant may be purged with very different efficiency depending on the ecological context in which selection acts.

The effect of competitive filtering on purging can also be tested empirically. Experimental populations could vary the number of candidates competing for a fixed number of breeding or recruitment positions while holding population size similar, then follow deleterious variants through time.

Genomic sampling before and after competitive recruitment could test whether deleterious alleles are preferentially removed before entering the breeding population. Long-term studies that combine genomic load with mating or reproductive success provide another test, as illustrated by the association between genomic load and male mating success in black grouse [13].

Nearly neutral theory describes the balance between selection and drift, but the strength with which a fitness difference is realised also depends on how individuals compete for survival and reproduction. Consequently, the evolutionary fate of weakly deleterious variation depends on both its genetic effect and the mode through which that effect is exposed to selection.

## Ethics

This study used simulations only and required no ethical approval.

## Data accessibility

Simulation scripts are available from GitHub at https://github.com/TBirley/SLiM_Selection_Mode_SFS. The Electronic Supplementary Material contains the full simulation methods and sensitivity analyses.

## Authors’ contributions

T.B. ran the simulations and generated the primary results. C.v.O. developed the conceptual framing and drafted the manuscript. Both authors revised and approved the final manuscript.

## Acknowledgements

We thank Hernán E. Morales, Jim J. Groombridge, Johanna Winder and Simone Immler for discussions about genetic load and sexual selection.

## References

1. Kimura M. 1968 Evolutionary rate at the molecular level. Nature 217, 624–626. 10.1038/217624a0

2. Ohta T. 1973 Slightly deleterious mutant substitutions in evolution. Nature 246, 96–98. 10.1038/246096a0

3. Charlesworth B. 2009 Effective population size and patterns of molecular evolution and variation. Nature Reviews Genetics 10, 195–205. 10.1038/nrg2526

4. de Jong MJ, van Oosterhout C, Hoelzel AR, Janke A. 2024 Moderating the neutralist-selectionist debate: exactly which propositions are we debating, and which arguments are valid? Biological Reviews 99, 23–55. 10.1111/brv.13010

5. Wallace B. 1975 Hard and soft selection revisited. Evolution 29, 465–473. 10.1111/j.1558-5646.1975.tb00836.x

6. Reznick D. 2016 Hard and soft selection revisited: how evolution by natural selection works in the real world. Journal of Heredity 107, 3–14. 10.1093/jhered/esv076

7. Bell DA, Kovach RP, Robinson ZL, Whiteley AR, Reed TE. 2021 The ecological causes and consequences of hard and soft selection. Ecology Letters 24, 1505–1521. 10.1111/ele.13754

8. Whitlock MC. 2000 Fixation of new alleles and the extinction of small populations: drift load, beneficial alleles, and sexual selection. Evolution 54, 1855–1861. 10.1111/j.0014-3820.2000.tb01232.x

9. Siller S. 2001 Sexual selection and the maintenance of sex. Nature 411, 689–692. 10.1038/35079578

10. Whitlock MC, Agrawal AF. 2009 Purging the genome with sexual selection: reducing mutation load through selection on males. Evolution 63, 569–582. 10.1111/j.1558-5646.2008.00558.x

11. Rowe L, Houle D. 1996 The lek paradox and the capture of genetic variance by condition dependent traits. Proceedings of the Royal Society B 263, 1415–1421. 10.1098/rspb.1996.0207

12. Wanders K, Chen G, Feng S, Zhang G, Székely T, Bruford M, Végvári Z, Eichhorn G, Urrutia A. 2023 Polygamy and purifying selection in birds. Evolution 77, 276–288. 10.1093/evolut/qpac010

13. Chen RS, et al. 2025 Predicted deleterious mutations reveal the genetic architecture of male reproductive success in a lekking bird. Nature Ecology & Evolution 9, 1924–1937. 10.1038/s41559-025-02802-8

14. Pointer MD, et al. 2026 Sexual selection purges mutation load, but not overall genetic diversity, decreasing vulnerability to extinction. Proceedings of the National Academy of Sciences USA 123, e2608675123. 10.1073/pnas.2608675123

15. Haller BC, Messer PW. 2023 SLiM 4: multispecies eco-evolutionary modeling. The American Naturalist 201, E127–E139. 10.1086/723601

16. Bertorelle G, et al. 2022 Genetic load: genomic estimates and applications in non-model animals. Nature Reviews Genetics 23, 492–503. 10.1038/s41576-022-00448-x

