## Supplementary material for "Selection mode changes the fate of nearly neutral deleterious mutations"

**Supplementary electronic material**

Forward-time simulations were conducted in SLiM 4.3.0 (Haller and Messer, 2023) using diploid, hermaphroditic populations with non-overlapping generations. Following selection, all surviving individuals mated once at random to produce the next generation. The genome consisted of 20 freely recombining autosomes. Each autosome contained 1,000 genes of 1,500 bp. Recombination occurred at a rate of 1 x 10^-9^ within genes and 1 x 10^-3^ between genes, while autosomes assorted independently. This genome architecture has been used extensively in previous SLiM studies (Kyriazis et al, 2021; Robinson et al., 2022; Al Hikmani et al., 2024). A focal uniform mutation rate of 1 x 10^-8^ was used.

Two modes of selection were examined: hard selection and soft selection. We followed Wallace’s (1975) framework, with definitions and biological examples summarised by Bell et al. (2021). Under hard selection, individual survival probability was equal to individual fitness, *w_i_*. If the number of survivors exceeded the carrying capacity, excess individuals were removed at random until population size equalled *K* = 500. Thus, in both selection treatments, 500 individuals contributed to the next generation after selection. Because all survivors mated once at random, the simulated breeding population had an intended *N_e_* ≅ 500. This makes the conventional codominant near-neutrality threshold |*s*| = 1/(2*N_e_*) = 0.001. In a separate analysis, we also explored mutations with selection coefficients that differed by one order of magnitude, i.e. *s* = 0.01 and *s* = 10^-4^.

Under soft selection, individuals competed directly for *K* survival opportunities, and we refer to this as our global rank-based model. Individuals were ranked by a competitive score equal to 0.5*w_i_* + U(0,0.5), where U(0,0.5) is a random draw from a uniform distribution. Individuals with the highest competitive scores survived. This approach was adopted because ranking individuals solely by *w_i_* produced near-deterministic selection. The soft-selection implementation was deliberately idealised to capture the core contrast between absolute filtering and relative competition. Thus, fecundity 50 produced many more candidates for the fixed $K=500$ opportunities than fecundity 5.

As a sensitivity analysis, we also implemented soft selection as repeated pairwise contests. The offspring cohort was randomly divided into two groups, and individuals from one group were paired against individuals from the other. Each individual received the same competitive score used in the global rank-based model, $0.5w_{i}+U(0,0.5)$. In each pair, the individual with the higher score survived to the next round; ties were resolved at random. Pairwise rounds continued until the number of survivors was $K$, or until fewer than $2K$ individuals remained. In the latter case, only the number of pairwise contests needed to reduce the population to $K$ was performed. Thus, for fecundity 5, the initial cohort of 2,500 individuals was reduced through 2,500 → 1,250 → 625 → 500, whereas for fecundity 50, the initial cohort of 25,000 individuals was reduced through 25,000 → 12,500 → 6,250 → 3,125 → 1,563 → 782 → 500.

To examine the effect of selection mode, simulations focused on two mutation classes: neutral mutations (*s* = 0) and weakly deleterious mutations (*s* = -0.001). Neutral mutations provided the baseline expectation for mutation accumulation in the absence of selection. Deleterious mutations had a dominance coefficient *h* = 0.01. Simulations were conducted under fecundities of 5, 20, and 50 offspring per breeding pair. Populations evolved for 5,000 generations, with mutation frequencies and mutation ages recorded every 100 generations. Each parameter combination was run for 100 replicate simulations. Mutation accumulation and site-frequency spectra (SFS) were compared among neutral evolution, hard selection and soft selection, and across fecundity treatments.

To examine whether treatments experienced comparable levels of drift, realised *N_e_* was estimated from selectively neutral marker mutations. Marker mutations were introduced during the 50,000-generation neutral burn-in at 2.5 × 10^−9^ mutations per site per generation and retained throughout the 5,000-generation experimental phase. These markers were used only for *N_e_* estimation. Focal mutations, used for mutation-accumulation, mutation-age and site-frequency-spectrum analyses, were introduced during the experimental phase at 1 × 10^−8^ mutations per site per generation. Realised *N_e_* was estimated using four complementary approaches: diversity-based *N_e_*, temporal *N_e_*, coalescent *N_e_* and reproductive-variance *N_e_*. These estimators capture different consequences of genetic drift and effective population size (Wang 2016).

Diversity-based *N_e_* was estimated from nucleotide diversity following Nei and Tajima (1981):

$$N_{e} = \frac{\pi}{4\mu}$$

where *μ* = 2.5 × 10^−9^ is the neutral-marker mutation rate.

Temporal *N_e_* was estimated from the standardised change in neutral allele frequencies between generations 4,900 and 5,000, following the temporal method of Waples (1989).

Coalescent *N_e_* was estimated from mean pairwise coalescence times in the SLiM tree-sequence outputs. Under the standard diploid neutral coalescent, the expected pairwise coalescence time is approximately 2*N_e_* generations (Hudson 1990), so:

$$N_{e} = \frac{T_{pair}}{2}$$

where *T_pair_* is the mean pairwise coalescence time in generations.

Reproductive-variance *N_e_* was estimated from the variance in realised reproductive success following Crow and Denniston (1988) as:

$$N_{e} = \frac{4N_{br} - 2}{V_{k} + 2}$$

where *N_br_* is the number of breeding individuals and *V_k_* is the variance in reproductive success.

Our simple model was designed to isolate the essential contrast between absolute fitness filtering and relative competitive filtering. We therefore focussed on a single weakly deleterious mutation class rather than a full distribution of fitness effects (DFE), and a simple high-discrimination ranking scheme rather than a species-specific competitive process. Future studies should explore more biologically realistic DFEs, dominance coefficients, spatial structure, different types of competition, different level of environmental variation, repeated life-stage filtering, and variation in the number of competitors and winners. At the same time, the genome architecture, mutation rate, recombination scheme and basic SLiM framework were kept close to previous applications of this model class to maintain comparability with earlier studies (Kyriazis et al., 2021; Robinson et al., 2022; Al Hikmani et al., 2024).

Simulation scripts are available at <https://github.com/TBirley/SLiM_Selection_Mode_SFS>.

**Results**

**Realised *N_e_***

Across diversity-, temporal-, coalescent- and reproductive-variance-based estimators, realised *Nₑ* remained close to the target population size (Table S1). Selection mode had little effect on realised *Nₑ*, indicating that differences in mutation accumulation and the site-frequency spectrum were not driven by systematic differences in genetic drift among selection treatments. The main source of variation was fecundity, with low-fecundity treatments producing slightly higher *Nₑ* estimates than high-fecundity treatments. This pattern is explained by lower variance in realised reproductive success (*V*_k_) in low-fecundity treatments: *V*_k_ was approximately 1.60 at fecundity 5, but close to the Wright-Fisher expectation of 2 at fecundity 50 (Table S2). Overall, these estimates confirm that *s* = −0.001 remained close to the classical 1/(2*Nₑ*) scale across treatments.

**Sensitivity to deleterious effect size under hard selection**

As a further check, we compared mutation accumulation under hard selection for deleterious-effect sizes below, near and above the intended nearly neutral threshold. Mutations with *s*=-10^-4^, an order of magnitude below the 1/(2*N_e_*) threshold, accumulated close to the neutral expectation (Figure S1). Mutations with *s*=-0.001, close to the nearly neutral threshold, were partially constrained under hard selection but continued to accumulate over time. Mutations with *s*=-0.01, an order of magnitude above the threshold, experienced efficient purifying selection. Fecundity had little qualitative effect under hard selection across these effect sizes, supporting the conclusion that increased fecundity alone does not greatly improve purging when selection acts through absolute survival probabilities.

**Pairwise-contest soft selection**

To test whether the effect of soft selection depended on global ranking, we repeated the simulations using a pairwise-contest implementation of soft selection. In this model, individuals competed in repeated one-versus-one contests until *K* survivors remained. The pairwise-contest model produced qualitatively similar results to the global rank-based model. At fecundity 50, both soft-selection implementations strongly reduced mutation accumulation relative to hard selection and neutral evolution (Figure S2a). Across fecundities, mutation accumulation declined as the number of candidates increased, and both soft-selection implementations showed the strongest filtering at fecundity 50 (Figure S2b). The pairwise-contest model was slightly less efficient than cohort-wide ranking, but the overall conclusion was unchanged: relative competition can strongly reduce the accumulation of weakly deleterious mutations, and the strength of this effect depends on the intensity of competitive filtering.

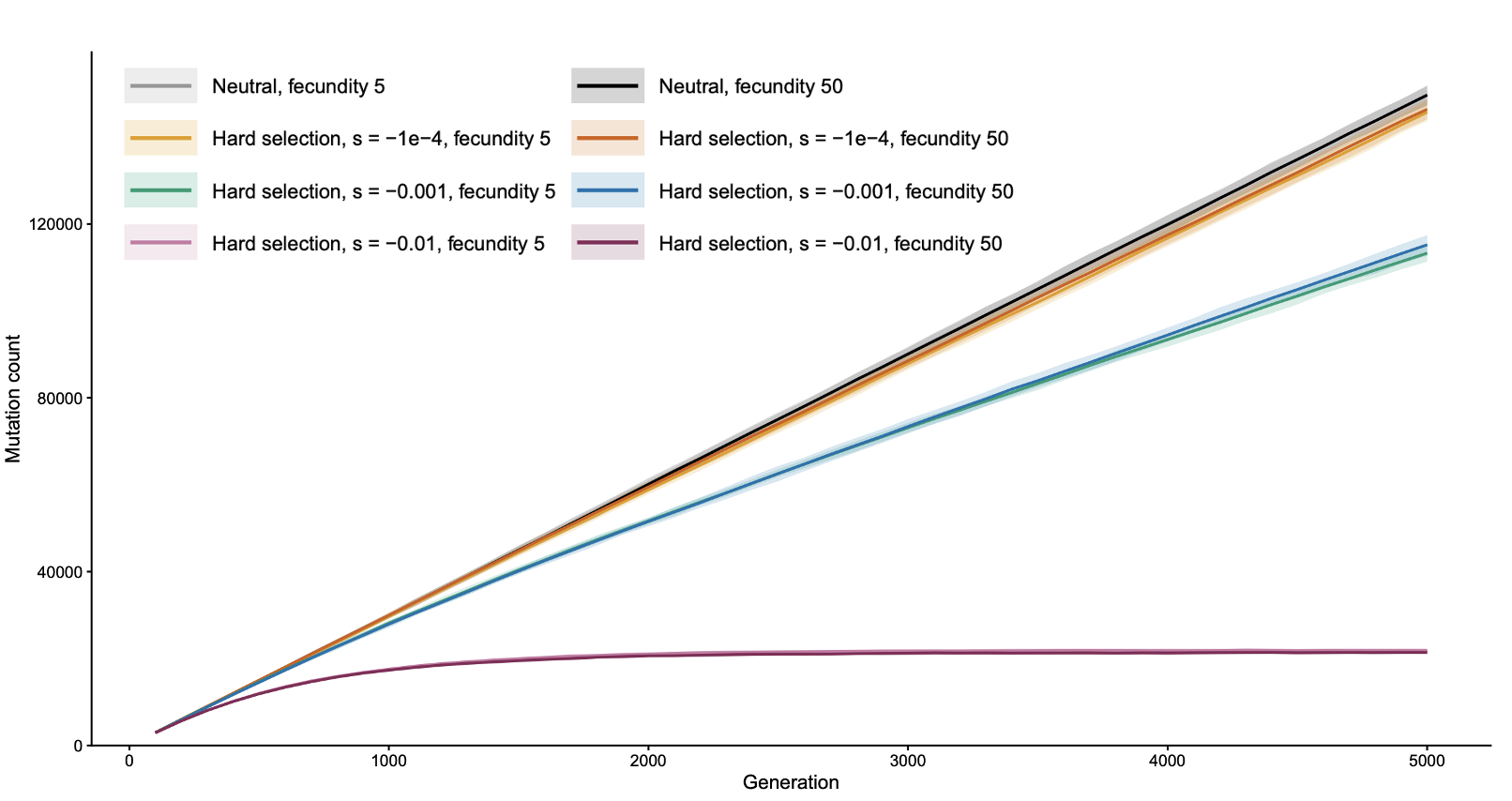

**Figure S1. Hard-selection sensitivity to selection coefficient.** Mutation accumulation over 5000 generations under neutral evolution and hard selection for mutations with *s*=-10^-4^, -0.001, and -0.01, with fecundity set to 5 or 50 offspring per breeding pair. These deleterious-effect sizes fall below, near and above the intended classical (1/(2*N_e_*)) threshold, respectively. Under hard selection, mutations below the threshold accumulated close to the neutral expectation, mutations near the threshold were partially constrained but continued to accumulate, and mutations above the threshold were strongly constrained. Fecundity had little qualitative effect on mutation accumulation under hard selection.

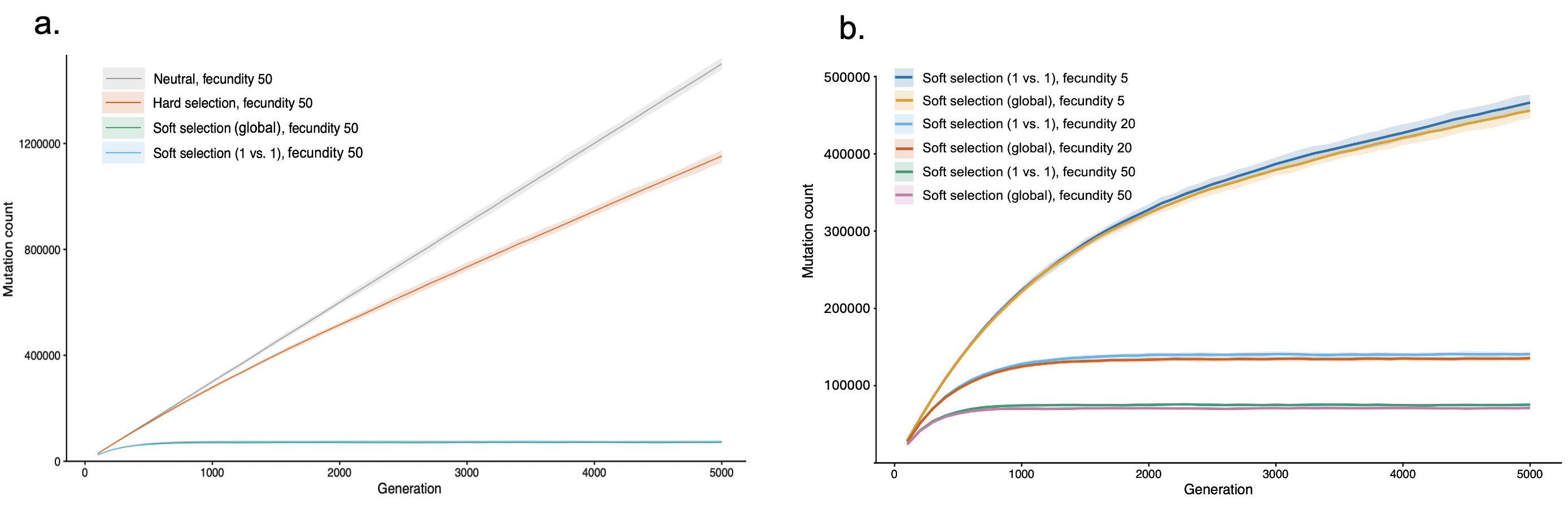

**Figure S2. Pairwise-contest soft selection produces qualitatively similar filtering to cohort-wide ranking.** (a) Accumulation of weakly deleterious mutations (*s* = -0.001) over 5,000 generations under neutral evolution, hard selection, global rank-based soft selection and pairwise-contest soft selection at fecundity 50. Both soft-selection implementations strongly reduce mutation accumulation relative to hard selection and neutral evolution. (b) Mutation accumulation under cohort-wide and pairwise-contest soft selection across fecundities 5, 20 and 50. Filtering is strongest at high fecundity, where more candidates compete for the same number of survival opportunities. Lines show mean mutation counts across replicate simulations; shaded bands show standard errors.

**Table S1. Realised effective population size estimates obtained from nucleotide diversity, temporal allele-frequency change, coalescent histories and reproductive variance.** Values are means ± SD across replicate simulations.

| **Selection mode** | **Neutral** | | **Hard selection** | | **Soft selection** | | |
| --- | --- | --- | --- | --- | --- | --- | --- |
| **Fecundity** | **5** | **50** | **5** | **50** | **5** | **20** | **50** |
| **Diversity-based *N_e_*** | 561.5 ± 14.6 | 510.7 ± 15.5 | 563.8 ± 14.8 | 514.2 ± 16.9 | 560.3 ± 16.4 | 518.8 ± 15.4 | 510.6 ± 15.0 |
| **Temporal *N_e_*** | 508.2 ± 23.2 | 482.6 ± 26.3 | 511.0 ± 22.4 | 475.1 ± 23.2 | 512.4 ± 24.9 | 486.3 ± 25.5 | 476.9 ± 26.5 |
| **Coalescent *N_e_*** | 554.8 ± 4.0 | 505.2 ± 3.0 | 557.7 ± 3.3 | 508.5 ± 3.5 | 561.0 ± 3.4 | 508.2 ± 2.8 | 495.0 ± 3.6 |
| **Reproductive- variance *N_e_*** | 556.1 ± 0.3 | 506.0 ± 0.3 | 556.1 ± 0.3 | 505.9 ± 0.3 | 555.8 ± 0.3 | 512.0 ± 0.3 | 502.6 ± 0.4 |

**Table S2. Variance in realised reproductive success (*V_k_*) across neutral, hard-selection and soft-selection simulations.** Values are mean *V*_k_ ± SD across replicate simulations. Low-fecundity treatments had lower variance in realised reproductive success, explaining why reproductive-variance *N*_e_ estimates were slightly above the census number of breeders.

| **Selection mode** | **Fecundity** | **Mean Vₖ** | **SD Vₖ** |
| --- | --- | --- | --- |
| Neutral | 5 | 1.60 | 0.0019 |
| Neutral | 50 | 1.96 | 0.0024 |
| Hard selection | 5 | 1.60 | 0.0019 |
| Hard selection | 50 | 1.96 | 0.0027 |
| Soft selection | 5 | 1.60 | 0.0019 |
| Soft selection | 20 | 1.91 | 0.0024 |
| Soft selection | 50 | 1.98 | 0.0029 |

**References**

Al Hikmani, H., van Oosterhout, C., Birley, T., Labisko, J., Jackson, H.A., Spalton, A., Tollington, S. and Groombridge, J.J., 2024. Can genetic rescue help save Arabia's last big cat? Evolutionary Applications, 17(5), p.e13701. <https://doi.org/10.1111/eva.13701>

Bell, D.A., Kovach, R.P., Robinson, Z.L., Whiteley, A.R. and Reed, T.E., 2021. The ecological causes and consequences of hard and soft selection. Ecology Letters, 24(7), pp.1505-1521. <https://doi.org/10.1111/ele.13754>

Crow JF, Denniston C. 1988 Inbreeding and variance effective population numbers. Evolution 42, 482–495. <https://doi.org/10.1111/j.1558-5646.1988.tb04154.x>

Haller BC, Messer PW. 2023 SLiM 4: multispecies eco-evolutionary modeling. The American Naturalist 201, E127-E139. <https://doi.org/10.1086/723601>

Hudson RR. 1990 Gene genealogies and the coalescent process. Oxford Surveys in Evolutionary Biology 7, 1–44.

Kyriazis, C.C., Wayne, R.K. and Lohmueller, K.E., 2021. Strongly deleterious mutations are a primary determinant of extinction risk due to inbreeding depression. Evolution Letters, 5(1), pp.33-47. <https://doi.org/10.1002/evl3.209>

Nei M, Tajima F. 1981 Genetic drift and estimation of effective population size. Genetics 98, 625–640. <https://doi.org/10.1093/genetics/98.3.625>

Robinson, J.A., Kyriazis, C.C., Nigenda-Morales, S.F., Beichman, A.C., Rojas-Bracho, L., Robertson, K.M., Fontaine, M.C., Wayne, R.K., Lohmueller, K.E., Taylor, B.L. and Morin, P.A., 2022. The critically endangered vaquita is not doomed to extinction by inbreeding depression. Science, 376(6593), pp.635-639. <https://doi.org/10.1126/science.abm1742>

Wallace, B., 1975. Hard and soft selection revisited. Evolution, pp.465-473. <https://doi.org/10.1111/j.1558-5646.1975.tb00836.x>

Wang J. 2016 Prediction and estimation of effective population size. Heredity 117, 193–206. <https://doi.org/10.1038/hdy.2016.43>

Waples RS. 1989 A generalized approach for estimating effective population size from temporal changes in allele frequency. Genetics 121, 379–391. <https://doi.org/10.1093/genetics/121.2.379>
